# Development of a Xylene-Free Sample Preparation Protocol for Quantitative Proteomics of Clinically Relevant Formaldehyde-Fixed Paraffin-Embedded Needle Biopsy Samples

**DOI:** 10.64898/2026.05.12.724492

**Authors:** Gontse Mabuse Moagi, Lívia Beke, Gábor Méhes, Gábor Kecskeméti, Zoltán Szabó, Lilla Turiak, Éva Csősz

## Abstract

Fresh-frozen tissues are considered the gold standard for proteomic analyses due to superior preservation of protein integrity; however, their use is limited by the logistical and financial requirements of long-term storage. Formaldehyde-fixed paraffin-embedded (FFPE) tissues provide a practical alternative owing to their stability and widespread availability in clinical settings. A critical step in FFPE proteomics is deparaffinization, which traditionally relies on organic solvents such as xylene, along with efficient reversal of formaldehyde-induced crosslinks. In this study, we evaluated multiple FFPE protein extraction and digestion workflows including chaotropic, surfactant-based, and detergent-free approaches in combination with xylene-free deparaffinization strategies, using label-free data-independent acquisition (DIA) LC-MS/MS. Among the tested methods, a chaotropic-, reductant-, and surfactant-free in-solution digestion workflow demonstrated robust protein and peptide recovery. A modified version of this protocol further improved peptide coverage while maintaining comparable protein depth.

The applicability of the optimized workflow was assessed using FFPE needle biopsy samples from control, hepatic steatosis, and liver fibrosis groups. Distinct proteomic patterns were observed across conditions, with hepatic steatosis associated with early activation of stress-response pathways, while fibrosis showed evidence suggesting altered lipid metabolism.

Overall, this study presents a simple, xylene-free, and MS-compatible workflow for FFPE proteomics that is suitable for low-input clinical samples and may support broader application of archival tissues in proteomic research.

## 1. Introduction

Mass spectrometry (MS)-based proteomics, particularly liquid chromatography-tandem mass spectrometry (LC-MS/MS), has become a cornerstone for the comprehensive characterization of biological systems [1], [2]. While global proteomic analyses of bodily fluids are widely used to investigate disease mechanisms and predict clinical outcomes, tissue-based proteomics provides essential spatial and cellular context that cannot be captured from biofluids alone [3], [4]. Tissue specimens are commonly preserved as either fresh-frozen or formaldehyde-fixed paraffin-embedded (FFPE) samples. Fresh-frozen tissues are considered the gold standard for proteomic analyses due to superior preservation of protein integrity; however, their use is constrained by the logistical and financial demands of long-term cold storage. In contrast, FFPE tissues represent a highly accessible and stable alternative, enabling room-temperature storage while preserving tissue architecture for histopathological assessment [5]. Although FFPE preservation can compromise quantitative reproducibility relative to fresh-frozen tissues, previous studies have demonstrated substantial overlap in protein identifications and gene ontology profiles between the two approaches [6], [7], [8] supporting the applicability of FFPE specimens for proteomic investigations.

Proteomic analysis of FFPE tissues can be performed using scrolls [9] or section-based sampling approaches, including laser capture microdissection (LCM) and manual macrodissection (MMD) [10]. Among these, MMD offers practical advantages by enabling spatially resolved sampling guided by histopathological features, while avoiding the technical complexity and labor-intensive nature of LCM [11]. Consequently, MMD has become a widely adopted strategy in routine FFPE proteomics workflows.

A critical step in FFPE sample preparation is deparaffinization, which has traditionally relied on organic solvents such as xylene, followed by graded ethanol washes for rehydration [12], [13]. Although effective, these methods involve hazardous reagents and multiple processing steps. Alternative strategies have therefore been introduced, including commercial reagents like SafeClear [14] and proprietary systems such as adaptive focused acoustics (AFA) [15]. Solvent-free or “green” deparaffinization approaches such as hot water-based methods [16], [17], projected hot air deparaffinization (PHAD) [18], and oven heating [19] are prospective approaches due to their simplicity and reduced chemical usage.

Following deparaffinization, FFPE proteomic analysis requires reversal of formaldehyde-induced crosslinks to facilitate efficient protein extraction and enzymatic digestion. These crosslinks, primarily methylene bridges, reduce protease accessibility, particularly for membrane-associated and low-abundance proteins, thereby limiting proteome coverage [20]. Antigen retrieval strategies typically combine heat with buffering systems and often include strong detergents such as sodium dodecyl sulfate (SDS) or sodium deoxycholate (SDC) to enhance protein solubilization. However, such detergents are incompatible with LC-MS/MS and require additional cleanup steps, including protein precipitation [10] or filter-based buffer exchange [21], which can be problematic for low-input samples and may result in sample loss.

In this study, we systematically evaluate multiple FFPE protein extraction and digestion workflows in combination with solvent-free deparaffinization strategies. By comparing chaotropic, surfactant-based, and detergent-free approaches—including a modified protocol optimized for low-input material - we aim to identify robust and MS-compatible methods that maximize proteome coverage. The optimized workflow was further applied to liver needle biopsy samples across progressive stages of liver damage, demonstrating its suitability for low-input proteomics and its potential to uncover biologically relevant proteomic alterations. This work provides a practical framework for improving FFPE-based proteomics, with an emphasis on relative simplicity, and compatibility with limited sample material.

## 2. Methods

### 2.1 Sample Information

FFPE sample material used for protocol optimization consisted of freely available archived patient tissue obtained from routine diagnostic procedures. For the disease cohort, inclusion criteria comprised patients with normal liver, fibrosis, or hepatic steatosis, aged between 32 and 72 years, with a balanced gender distribution of 1:1 female to male.

Needle biopsy specimens were fixed in 4% paraformaldehyde at 4 °C for 6 h, followed by sequential dehydration in 75%, 80%, 90%, 95%, and 100% ethanol. The resulting FFPE blocks were sectioned at a thickness of 10 μm using a HM34E rotation microtome (Epredia Hungary, Budapest) according to standard protocols and mounted onto Superfrost™ glass slides. This work was approved by the Scientific and Research Ethics Committee of the Scientific Council of Health of the Hungarian Government under the registration number of IV/8465-3/2021/EKU.

### 2.2 Protein extraction from FFPE Tissue sections

All reagents used in the procedure were purchased from Sigma, unless otherwise specified. Serial formaldehyde-fixed paraffin-embedded (FFPE) tissue sections (10 μm thickness) were baked at 60 °C for 2 h to prevent tissue detachment. Deparaffinization was subsequently performed using either projected hot air deparaffinization (PHAD) or hot distilled water. For PHAD, a conventional hair dryer was used at maximum power for 20 min to melt paraffin. Slides were immediately immersed in distilled water for 5 min to facilitate rehydration and solidification of residual paraffin. For the hot water method, 1 mL of distilled water at 80 °C was applied dropwise to melt and remove paraffin. A ruler was used to delineate the tissue area for consistent sampling and the tissue samples were collected with the help of a sterile scalpel.

The resulting deparaffinized tissue slices were subjected to further processing and digestion according to the following protocols.

#### 2.2.1 Approach 1: Chaotropic-Based in-solution digestion (App 1)

Heat-induced antigen retrieval (HIAR) was performed prior to tissue collection by immersing slides in citrate buffer (95 mM trisodium citrate, 21 mM citric acid, pH 6.0) at 75 °C for 60 min. Tissue was then scraped off, resuspended in 75 μL of 8 M urea, and solubilized by 15 cycles of sonication (1 min on, 30 s off) in a chilled (∼1.5 °C) water bath. Samples were reduced with 10 mM dithiothreitol (DTT) for 30 min at 37 °C and alkylated with 15 mM iodoacetamide (IAA) for 20 min at room temperature in the dark. The solution was diluted to 4 M urea with 50 mM Tris-HCl (pH 8.0) and incubated with 40 ng Pierce Trypsin/Lys-C (ThermoFisher Scientific) for 4 h at 37 °C. Urea concentration was further reduced to 0.8 M prior to overnight digestion with 60 ng trypsin at 37 °C. Digestion was terminated by acidification with 6% trifluoroacetic acid (TFA) to pH < 3.

#### 2.2.2 Approach 2: Chaotropic-, Reductant-, and Surfactant-Free in-solution digestion (App_2)

This approach was adapted from a previously described method [22]. Collected tissue was washed with 50 μL of 100% acetonitrile (ACN) and dried using a SpeedVac (ThermoFisher Scientific). The dried sample was resuspended in 40 μL of 60 mM triethylammonium bicarbonate (TEAB) and incubated at 95 °C for 60 min. Subsequently, 10 μL of 60% ACN in 60 mM TEAB was added, followed by incubation at 75 °C for an additional 60 min. Pre-digestion was performed with 40 ng Trypsin/Lys-C for 4 h at 37 °C, followed by overnight digestion with 60 ng trypsin (16 h, 37 °C). The reaction was terminated by acidification with 6% TFA to pH < 3.

#### 2.2.3 Approach 3: On-Surface Surfactant-Based in-solution digestion (App_3)

This protocol was adapted with minor modifications [10]. Tissue was solubilized in 100 μL of freshly prepared buffer (80 mM HEPES, pH 8.0, 80 mM DTT, 4% SDS). Samples underwent two rounds of sonication (each round consisted of 15 cycles: 1 min on, 30 s off) in a chilled (∼1.5 °C) water bath, followed by incubation at 99 °C for 1 h. After centrifugation (15 min, 13,000 × g, RT), the supernatant was collected and alkylated with IAA to a final concentration of 15 mM for 30 min in the dark. Proteins were precipitated with four volumes of ice-cold acetone and incubated overnight at −20 °C. Samples were centrifuged (40 min, 16,000 × g, 4 °C), and pellets were washed twice with 80% ice-cold acetone. Pellets were resuspended in 25 μL digestion buffer (1.5 M urea, 100 mM HEPES, pH 8.0) using sonication (3 cycles). Trypsin/Lys-C digestion (1:30, w/w) was performed for 4 h at 37 °C (600 rpm), followed by dilution to 0.75 M urea and overnight trypsin digestion (1:20, w/w). Digestion was quenched with TFA (pH < 3).

#### 2.2.4 Approach 4: On-Surface Surfactant-Based digestion (App 4)

This method was adapted with minor modifications [23]. HIAR was performed as described in App 1. Reduction was carried out by applying a 3 μL droplet (20% glycerol, 0.1% SDS, 5 mM DTT) and incubating for 40 min at 37 °C. Alkylation was performed using a 3 μL droplet containing 10 mM IAA in 25 mM ammonium bicarbonate (ABC) for 20 min at room temperature in the dark. Pre-digestion was conducted in two cycles using 3 μL droplets (20% glycerol, 40 ng Trypsin/Lys-C in 50 mM ABC) for 40 min at 37 °C in a humidified chamber. Final digestion was performed in three cycles with 200 ng trypsin under the same conditions. Peptides were extracted by repeated pipetting (5×) with 3 μL of 10% acetic acid and collected into LoBind tubes.

#### 2.2.5 Approach 2M: Chaotropic- and Surfactant-Free Buffer with Reduction/Alkylation (App_2M)

This modified protocol incorporated reduction and alkylation into App_2. Following incubation at 95 °C in 60 mM TEAB, samples were reduced using 10 μL of 60% ACN containing 80 mM DTT and incubated at 75 °C for 60 min. Alkylation was performed with 15 mM IAA for 30 min at room temperature in the dark. Pre-digestion (40 ng Trypsin/Lys-C, 4 h at 37 °C) and subsequent steps were identical to those described for App_2.

Following all protocols, peptides were purified using C18 PierceTips (ThermoFisher Scientific), dried, and stored at -20 °C until further analysis.

### 2.3 Equal Loading of Samples

Peptide concentrations were estimated by measuring absorbance at 280 nm (A280) using a NanoDrop ND-1000 spectrophotometer (ThermoFisher Scientific), with the conversion factor set to “1 Abs = 1 mg/mL”. Samples were normalized to a final concentration of 80 ng/μL in 1% formic acid (FA) prior to analysis.

### 2.4 Chromatographic and Mass Spectrometric Condition

Digested samples were redissolved in 10 µL 1% FA and analyzed as single shot DIA analyses. For method optimization the LC-MS analysis was carried out on an EASY nLC-1200 coupled to Orbitrap Fusion (both from ThermoFisher Scientific). 5 µL of sample was injected at a constant flow of 1 µL/min solvent A to a 5 µm, 180 µm × 20mm NANOACUITY™ trap column packed with ACUITY UPLC™ Symmetry C18 particles (Waters). Then, analytical nanoflow separations were achieved using a 1.7 µm, 75 µm × 150mm ACUITY UPLC™ M-Class Peptide BEH C18 column (Waters). Peptides were eluted at a flow rate of 300 nL/min at a gradient of 0-2% solvent B over 2 min, followed by a rise to 25% of solvent B over 52 min and then to 38% solvent B over 11 min. Thereafter, solvent B was increased to 90% over 4 min and held for 3 min, after which the system returned to 2% solvent B in 1 min for a 2 min hold-on. For mass spectrometry analyses the nanoelectrospray ion source (NSI) (Nanospray Flex™, ThermoFisher Scientific) was operated at 2.3 kV in the positive-ion mode, with an ion-transfer tube set to 275 °C. Following MS1 spectra collection at 120,000 resolution in the range of 350-1200 m/z, precursor ions were fragmented at fixed isolation windows of 12 m/z with 55 scan events, in DIA mode, with HCD fragmentation using a 30% normalized collision energy to produce corresponding MS2 spectra. Product ions were detected in the Orbitrap analyzer with an AGC target of 2.0e5 and a maximum injection time of 60 ms, 30,000 resolutions in the range of 145-1,450 m/z.

Case study LC-MS analysis was carried out on a Waters ACQUITY UPLC M-Class LC system (Waters, Milford, MA, USA) coupled to Orbitrap Exploris™ 240 Mass Spectrometer (ThermoFisher Scientific). 2 µL of sample was injected at a constant flow of 7 µL/min solvent A to a Symmetry^®^ C18 (100 Å, 5 µm, 180 µm × 20 mm) trap column (Waters, Milford, MA, USA). Then, analytical nanoflow separations were achieved using an ACQUITY UPLC^®^ M-Class Peptide BEH C18 analytical column (130 Å, 1.7 µm, 75 µm × 250 mm). Peptides were eluted at a flow rate of 200 nL/min at a gradient of 3-8% solvent B over 1 min, followed by a rise to 32% of solvent B over 81 min and then to 40% solvent B over 3 min. Thereafter, solvent B was increased to 90% over 1 min and held for 2 min, after which the system returned to 3% solvent B in 1 min for a 15 min hold-on. For mass spectrometry analyses the NSI (Nanospray Flex™, ThermoFisher Scientific) was operated at 1.8 kV in the positive-ion mode, with an ion-transfer tube set to 280 °C. Following MS1 spectra collection at 30,000 resolution in the range of 380-985 m/z, precursor ions were fragmented at fixed isolation windows of 10 m/z with 60 scan events, in DIA mode, with HCD fragmentation using a 28% normalized collision energy to produce corresponding MS2 spectra. Product ions were detected in the Orbitrap analyzer with an AGC target of 2.0e6 and a maximum injection time of 40 ms, 11,250 resolutions and the DIA m/z range was set to auto.

### 2.5 Data Analysis

Raw files were processed using DIA-NN (version 2.2) [24], [25] in library-free search mode (Table S1). This tool supports endogenous peptides-based retention time alignment, as such, no indexed retention time (iRT) peptides were added in the samples. A human protein sequence FASTA database (20420 reviewed entries) was downloaded from UniProtKB on 2025-07-26, and ‘FASTA digest’ and ‘Deep-learning spectra’ were enabled for *in silico* library generation. Other parameters included: Trypsin/P with maximum 2 missed cleavage; 3 maximum number of variable modifications; protein N-terminal M excision on; C-Carbamidomethyl was set as fixed modification for all approaches but App_2, while M oxidation, and N-terminal acetylation were set as variable modification. The XICs output was enable and all other parameters were kept as default.

The DIA-NN PARQUET output files were imported into the DEF workflow [26] together with the corresponding XICs report and report.pg_matrix files. Quantification parameters were maintained at default settings, and the HaoGroup contaminant library [27] was applied. Protein quantities were defined using the DIA-NN PG.MaxLFQ algorithms.

To improve quantification reliability, proteins were required to be supported by at least two peptides, thereby excluding single-hit identifications. Peptides were retained only if they contained a minimum of four consecutive b- or y-ions. Furthermore, proteins were required to be identified in at least two out of three biological replicates to minimize false-negative identifications.

Statistical comparisons between groups for experiment 2 were performed using an unpaired t-test with Welch’s correction in GraphPad Prism (version 8.0.1 for Windows, GraphPad Software, San Diego, California USA, www.graphpad.com, accessed on 11/04/2026). Results were considered significant at p < 0.05.

Principal component analysis (PCA) and protein ID overlap (%) calculations were performed in python (version 3.12.7) using the sklearn.decomposition and pandas libraries, respectively. Venn diagrams were generated using the IntersectMe web application (version 9.7.0).

For the quantitative analysis we applied our optimized digestion method for the examination of liver needle biopsies, where the group-level contrasts were defined within the DEF “Group-Visuals” module [26] to enable direct comparison between experimental conditions. Three comparisons (control, liver fibrosis, and hepatic steatosis) were specified, allowing a maximum of one missing value per replicate. The resulting datasets were exported for analysis in MetaboAnalyst 5 [28]. Given the low-input nature of FFPE needle biopsy samples, missing values were expected, particularly for low-abundance proteins. Data were filtered using a low-variance filter, and missing values were imputed using a left-censored approach by replacing them with one-fifth of the minimum positive intensity value. Subsequently, data were median-normalized and log10-transformed.

Normalized abundances were used for unpaired two-tailed t-tests (95% confidence interval). Differential abundant analysis was performed on the transformed, normalized data with multiple hypothesis correction using the Benjamini-Krieger false discovery rate (FDR 1%). Proteins with q-values < 0.01 and absolute log2 fold changes > 1 were considered as differentially abundant proteins (DAPs).

DAPs were exported from MetaboAnalyst and tertiary analysis was performed using the Ingenuity Pathway Analysis (IPA) platform (QIAGEN Inc., Redwood City, CA, USA) to obtain functional annotations, protein interaction networks, canonical pathways, and associated biological processes. Pathway enrichment significance was evaluated using right-tailed Fisher’s exact tests, whereas activation or inhibition states were predicted using IPA-generated z-scores under default parameters. Upstream regulators, downstream biological processes, and their predicted causal interactions were identified using the Ingenuity Knowledge Base, with z-scores calculated according to the directional abundance changes of DAPs.

## 3. Results and Discussion

Optimization experiments were conducted in three consecutive steps. First, the sample preparation efficiency and was evaluated from a limited tissue area (∼0.025mm^3^) using four FFPE processing approaches (App 1-4). This was followed by the application of the best two approaches (App_2 and App_3) to assess the qualitative and quantitative differences between the results obtained by the application of the two selected approaches. Based on the results App_2 was further optimized and in the final step, the modified form of App_2, the App_2M was used in a case study to examine biological differences characteristic to different forms of liver disease.

### 3.1 Xylene-free methods of FFPE sample preparation for proteomics analyses

Our goal was to develop a simpler, safer, and economically feasible protocol without the need of application of often expensive instrumentation. While deparaffinization by water has been employed in proteomics studies [16], [17], PHAD is yet to gain proteomics utility. Using two types of xylene-free deparaffinization, various protocols were evaluated for deep coverage of the FFPE tissue proteome in an effort to maximize the number of protein groups (PGs) obtained given the limited tissue sample availability (approximately 0.025mm^3^) in the case of needle biopsies. Approaches 1-4 include a wide range of widely applied non-commercial techniques for reversing the cross-linked nature of FFPE tissues including heat-induced antigen retrieval (HAIR) in citrate, TEAB, and SDS containing buffers. Based on the number of peptides (Figure S1A) and PGs (Figure S1B) identified, App_2 (TEAB) and App_3 (SDS-based) resulted in the highest numbers and were chosen for further experiments.

#### 3.1.1 Method refinement

Three randomly selected regions from two consecutive needle biopsy slides were digested with App_2 and App_3, respectively, providing 3 biological replicates in each case. Proteomics data were processed in DEF and the peptide and protein lists were evaluated (Table S2). The number of PGs from App_2 and App_3 differed significantly (App_2 939 PGs, App_3 492 PGs) (Figure 2A). A substantial proportion of proteins was consistently detected across all replicates, while only a subset was identified in individual samples, (Figure 2B, 2C, S2), highlighting the technical robustness of App_2 and App_3. Studies have shown that using dual-protease with trypsin and Lys-C has a positive effect on proteome coverage [29], as such trypsin/Lys-C mix was used for all digestion methods, in order to maximize coverage and have a better comparability. While in more than 88% of peptides 0 missed cleavage was observed in both approaches, App_3 showed a 0 missed cleavage rate of approximately 6% higher (Figure 2D). This increased cleavage efficiency could be due to extensive reduction and homogenization with sonication in HEPES buffer containing DTT and SDS. This was also evident at physicochemical level, with App_3 showing an increased proportion of shorter peptides (7-12 amino acids) (Figure 2E), which suggests enhanced proteolytic cleavage efficiency. Nonetheless the distribution of proteins among subcellular locations was nearly identical (Figure 2F), suggesting that the two approaches were not biased towards any specific cellular compartment (Figure S2).

**Figure 1:**
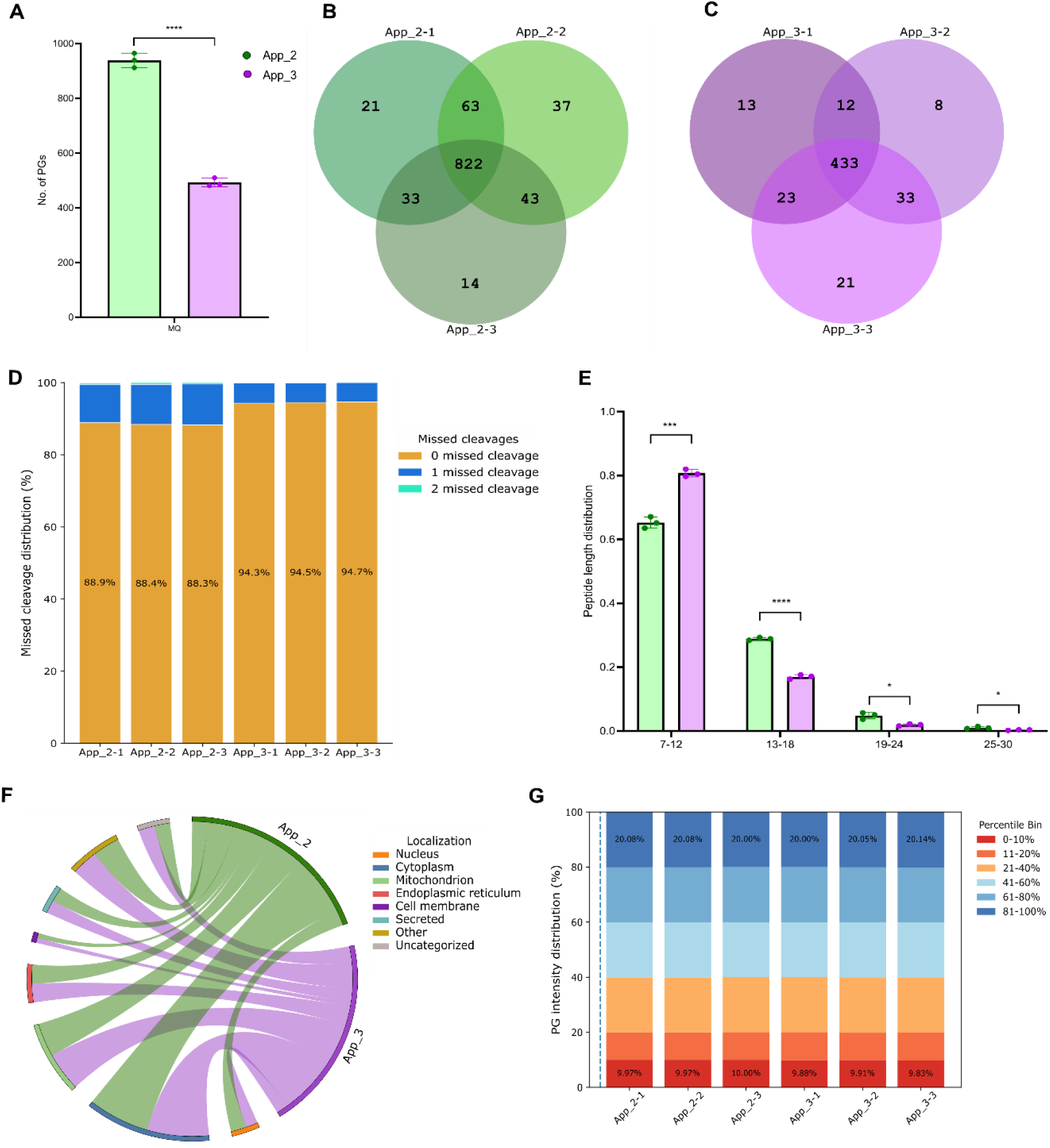
Comparative evaluation of Approach 2 and Approach 3 for FFPE proteomic analysis. (A) Comparison of the number of protein groups (PGs) identified using Approach 2 (App_2) and Approach 3 (App_3). Statistical significance was assessed using an unpaired t-test with Welch’s correction (**** p < 0.0001). (B, C) Venn diagrams of PG overlap across three technical replicates for (B) App_2 and (C) App_3. (D) Distribution of identified peptides according to the number of missed cleavages (0-2), reflecting digestion efficiency across both approaches. (E) Peptide length distribution presented as the number of identified peptides within defined amino acid length bins (7-12, 13-18, 19-24, and 25-30 residues). (F) The distribution of identified proteins across subcellular compartments for App_2 and App_3. (G) Distribution of PG intensities across percentile bins (0-10, 11-20, 21-40, 41-60, 61-80, and 81-100), highlighting detection sensitivity between the two approaches. “-1, -2, or -3” in the name of App_2 and App_3, respectively, refer to the technical replicates.

**Figure 2.**
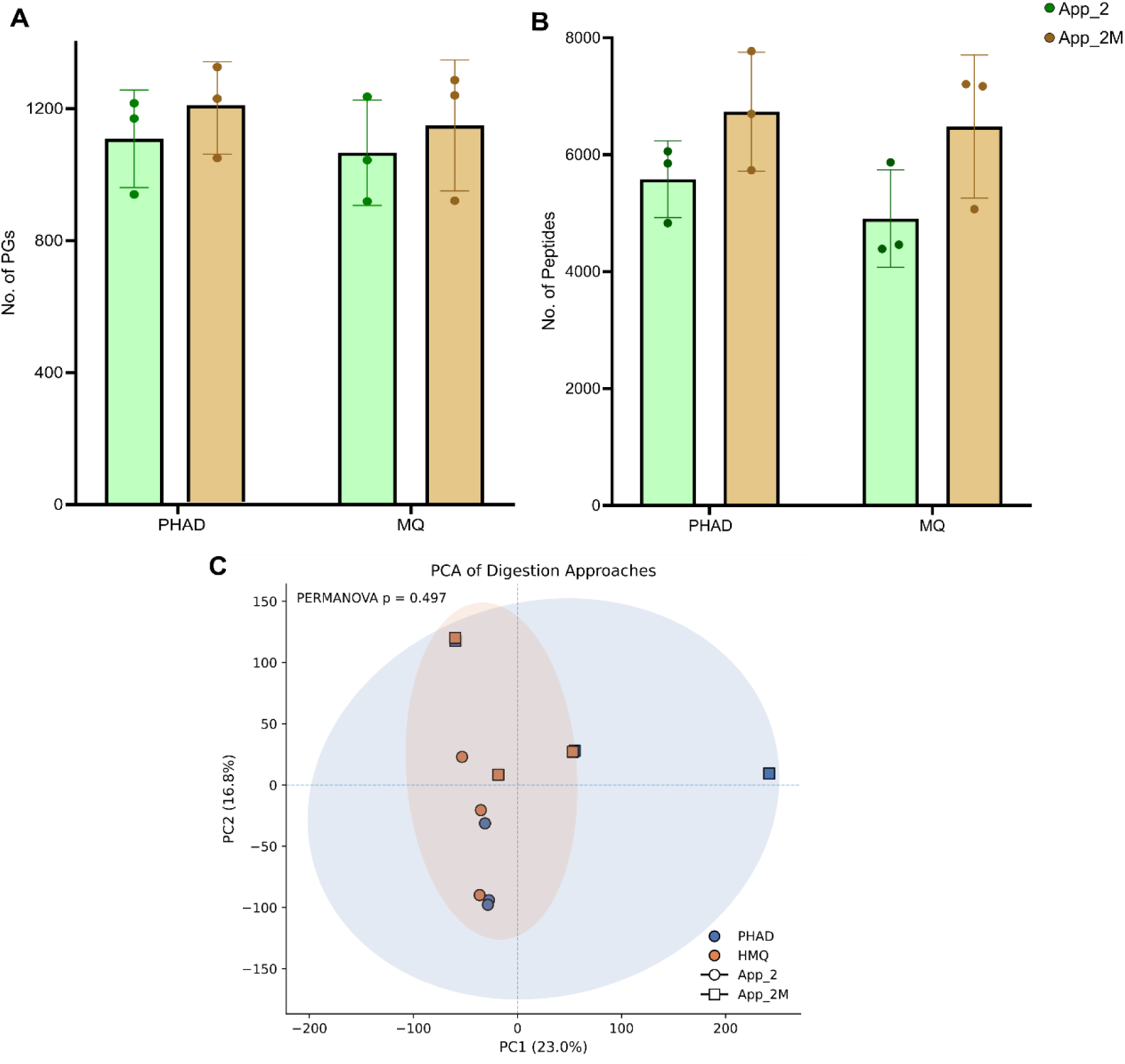
Comparative analysis of App_2 and App_2M using different deparaffinization methods. App_2 and App_2M were evaluated following deparaffinization with a projected hot air deparaffinization (PHAD) or hot Milli-Q water (HMQ). (A) Mean number of identified peptides and (B) protein groups (PGs), presented with standard deviation. (C) Principal component analysis of protein intensities derived from three replicates per approach and deparaffinization condition.

The distribution of quantified proteins across intensity percentiles revealed highly similar quantitative dynamic ranges (Figure 2G). Both approaches demonstrated comparable proportion of proteins in lower and higher percentile bins compared. This indicates that App2 and App_3 provide similar sensitivity and proteome depth.

Considering the number of posttranslationally modified peptides, no significant difference could be observed between App_2 and App_3 (Figure S3), indicating their similar performance in proteoform identification.

As App_2 provides more PGs involving less steps and it is easier to perform, we chose App_2 for further experiments. Based on the better cleavage efficiency observed in App_3, in the next step we modified App_2 by including reduction and alkylation as in App_3. The number of identified PGs showed a modest increase with the App_2M protocol compared to App_2 (Table S3). Mean PG counts were 1108 and 1202 for PHAD and 1066 and 1149 for HMQ-based deparaffinization, respectively. However, these differences did not reach statistical significance (Figure 3A). A similar pattern was seen with the number of identified peptides (Figure 3B). The principal component analysis of protein intensities from App_2 and App_2M showed no distinct clusters (Figure 3C), additionally, there was no significance in either the use of a hairdryer or deparaffinization with hot water.

**Figure 3.**
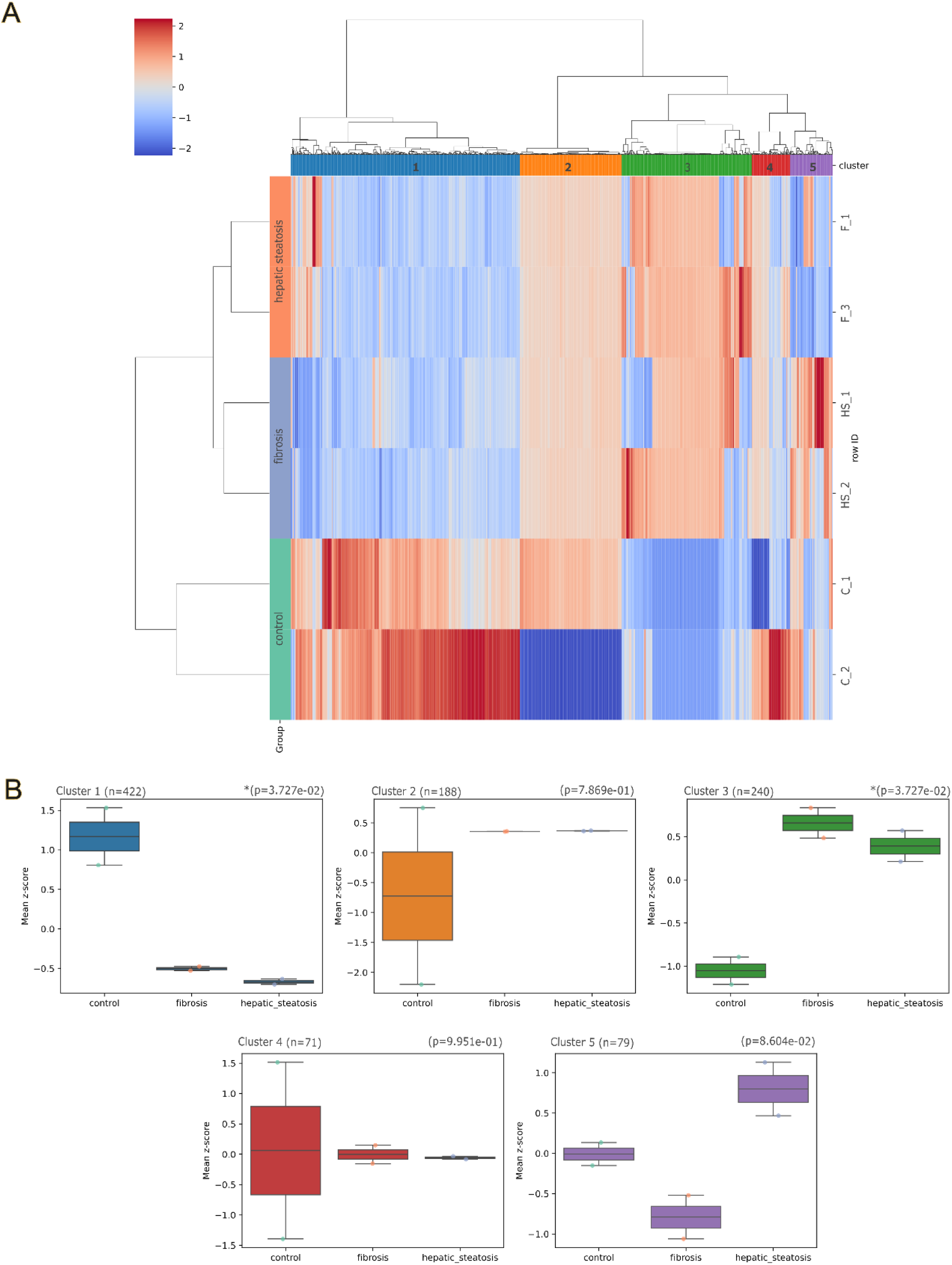
Protein abundance patterns across control, hepatic steatosis, and liver fibrosis groups. (A) Hierarchically clustered heat map of protein abundances across the three conditions. Proteins are grouped into five clusters (Clusters 1-5) based on similarity in expression profiles. Values represent Z-score-normalized intensities. (B) Box plots showing the distribution of mean Z-scores for each protein cluster (Clusters 1-5) across the three conditions.

We further evaluated a peptide to protein distribution to highlight approach depth (Table 1).

**Table 1.** Percentage of proteins identified by unique peptides. The columns shows the percentage of proteins identified by the number of unique peptides defined in the rows in case of different methods. Differences were calculated based on the comparison between App_2M and App_2. PHAD: projected hot air deparaffinization, HMQ: deparaffinization with hot MilliQ water

| Number of unique peptides per protein | PHAD-App_2 | PHAD-App_2M | Change | HMQ-App_2 | HMQ-App_2M | Change |
| --- | --- | --- | --- | --- | --- | --- |
| **1** | 40.88% | 37.89% | -2.99% | 42.48% | 37.76% | -4.72% |
| **2** | 17.57% | 17.25% | -0.32% | 17.80% | 16.92% | -0.88% |
| **3** | 10.08% | 10.15% | 0.07% | 10.39% | 10.88% | 0.49% |
| **4** | 6.59% | 7.30% | 0.71% | 7.41% | 6.72% | -0.69% |
| **5** | 5.05% | 5.59% | 0.54% | 4.47% | 4.68% | 0.21% |
| **6-20** | 17.06% | 18.73% | 1.67% | 15.45% | 19.62% | 4.17% |
| **>20** | 2.77% | 3.09% | 0.32% | 2.00% | 3.41% | 1.41% |

Although the dependence of PG identification on sequence counts is comparable across methods (Figure S4), App_2M showed an enrichment of protein groups supported by multiple unique peptides, particularly for proteins identified by 6-20 unique peptides. This change was significant particularly for HMQ-based deparaffinization, indicating improved peptide coverage per protein group (Figure S4). Our results show that modifying App_2 produced a protocol that delivers strong performance with fewer steps than the other approaches, resulting in App_2M that ultimately outperforms all previous protocols.

Extraction buffers containing chaotropes or detergents are known to interfere with the protease activity of trypsin [30] which results in incomplete protein digestion and consequently in lower proteome coverage due to oversampling of different cleavage forms of abundant peptides. However, with the inclusion of cleanup steps, surfactant-containing extraction buffers enhance trypsin activity and increase digestion efficiency [29], [31], [32]. This could be seen in the analysis of App 3 missed cleavage frequencies. The resultant low number of peptides could be due the loss during acetone precipitation. This suggests that App_3 could benefit from further refinements. Although considering time and simplicity, the improvement of App_2 was in line with our objective on a simple and user-friendly protocol.

Our results indicate that both App_2 and App_2M have good performance with respect to proteome integrity of the extracted proteins. They also show consistent protein abundances between technical replicates and a low percentage of miss cleaved peptides, possibly due to the use of a four-hour Lys-C predigestion with prior to overnight digestion with trypsin [33]. There is however a slight improvement in terms of number of quantified peptides and proteins and sequence coverage.

With our modified approach, we could identify about 1200 proteins which is consistent with previous reports of low-input FFPE proteomics using data-independent acquisition. Studies employing biopsy-scale or laser microdissected FFPE samples typically report proteome depths in the range of ∼800-2,000 proteins [15], [17], [34], reflecting the strong influence of sample input on proteome coverage.

### 3.2 Case study - examination of liver needle biopsy samples

Three biological replicates of FFPE needle biopsy samples (∼0.03-0.05 mm³) were analyzed, with App_2M, for each liver condition (fibrosis and hepatic steatosis), together with matched controls. The total number of identified proteins and peptides is summarized in Table 2, while detailed lists of identified peptides and proteins are provided in Table S4 and Table S5, respectively. Differences in the number of identified protein groups between control and disease samples may reflect both biological heterogeneity and variable protein recovery inherent to low-input FFPE proteomics workflows.

**Table 2:** Summary of the number of proteins and peptide identifications.

| Sample | No. of Peptides<br>(1% FDR) | No. of PGs |
| --- | --- | --- |
| LF-1 | 26416 | 2590 |
| LF-2 | 2931 | 452 |
| LF-3 | 26266 | 2583 |
| SH-1 | 25593 | 2583 |
| SH-2 | 26609 | 2589 |
| SH-3 | 2470 | 422 |
| C-1 | 11360 | 1161 |
| C-2 | 7708 | 1113 |
| C-3 | 571 | 83 |

Prior to downstream analysis, three samples (one from each condition) were excluded due to poor data quality. These samples exhibited a markedly reduced number of identified proteins and peptides, along with a disproportionately high contribution of hemoglobin and keratin (>50% of total signal), indicative of contamination (Figure S5). Consistently, these samples appear as outliers, primarily for the hepatic steatosis and fibrosis group, in principal component analysis (PCA) and heat maps (Figure S6).

Relative protein abundance patterns were visualized using a clustered heat map, revealing clear condition-dependent segregation (Figure 3A). Proteins with similar expression profiles were grouped into five distinct clusters (Figure 3B).

Differential abundance analysis using the *limma* model identified proteins with significant changes between conditions. Applying a log2 fold-change (log2FC) threshold of ≥1.5 and a Benjamini-Hochberg false discovery rate (FDR) q-value < 0.05, multiple proteins were found to be statistically differentially abundant across the disease states (Table S6). In comparison to controls, the hepatic steatosis group exhibited 11 proteins with decreased abundance and 147 proteins with increased abundance (Figure 4A). In fibrosis relative to controls, 23 proteins were found to be increased, while 151 were decreased in abundance (Figure 4B). When comparing hepatic steatosis to fibrosis, 11 proteins showed reduced and 13 increased abundances (Figure 4C).

**Figure 4.**
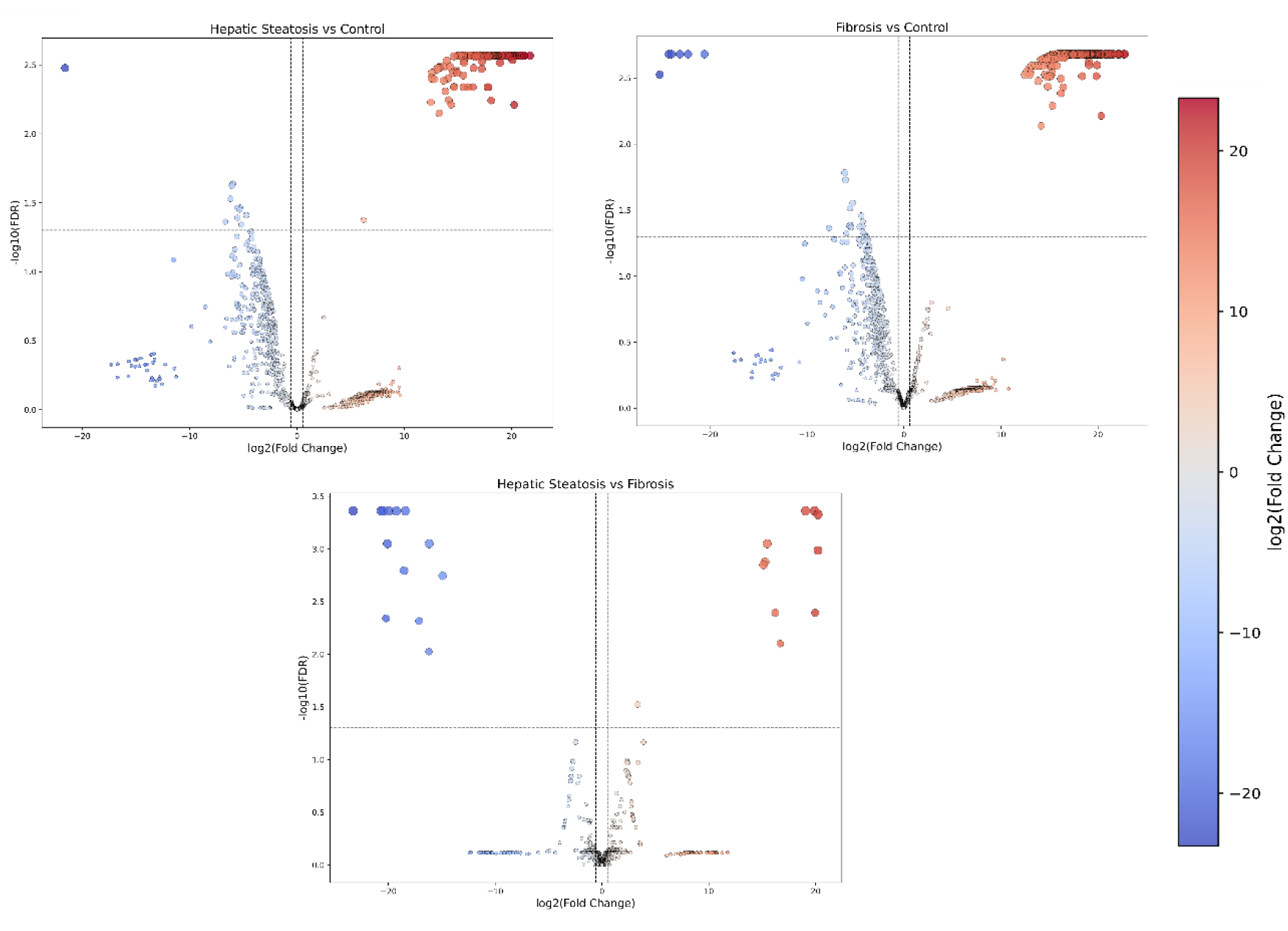
Differentially abundant proteins identified by *limma* analysis. (A-C) Volcano plots showing differentially abundant proteins between (A) hepatic steatosis and control, (B) liver fibrosis and control, and (C) hepatic steatosis and liver fibrosis. Differential abundance was defined using a log2 fold-change (log2FC) threshold of ≥1.5 and a Benjamini-Hochberg false discovery rate (FDR) q-value < 0.05. Proteins with increased abundance are indicated in red, whereas proteins with decreased abundance are shown in blue.

The list of differentially abundant proteins (DAPs) was further interrogated using regulator-effector network analysis in Ingenuity Pathway Analysis (IPA). Networks were generated separately for hepatic steatosis versus control (Figure 5A) and fibrosis versus control (Figure 5B) using the IPA Core Analysis with default settings. Predicted upstream regulators, downstream biological functions, and their causal relationships were inferred based on the Ingenuity Knowledge Base, with activation states assigned using z-scores derived from the directionality of protein abundance changes. Detail lists of all molecules and their respective relationships are provided in Table S7.

**Figure 5:**
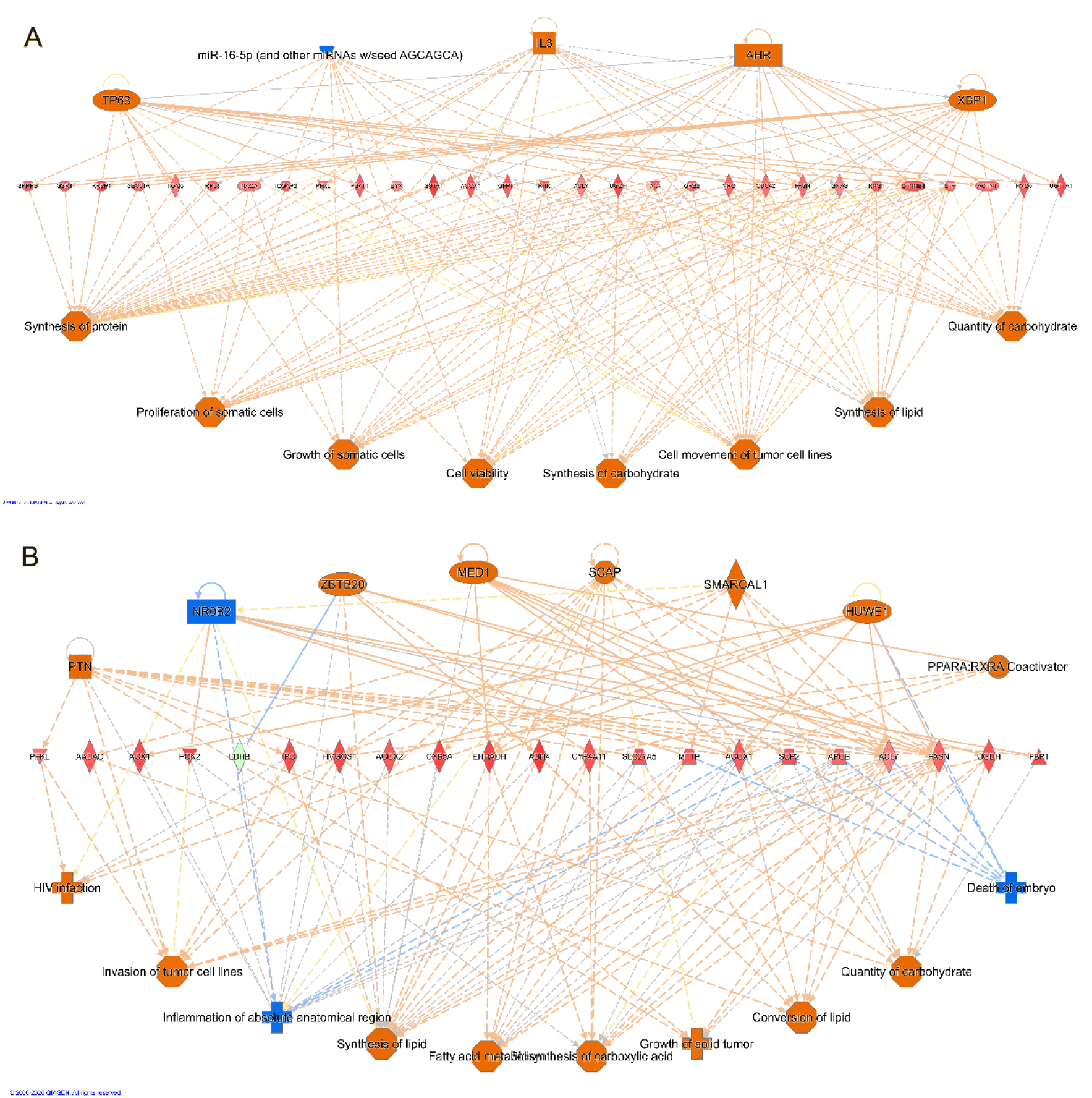
IPA regulator-effector networks of differentially abundant proteins. Regulator-effector networks generated by Ingenuity Pathway Analysis (IPA) for hepatic steatosis versus control (A) and fibrosis versus control (B). Node color reflects predicted activity state, with red indicating activation and blue indicating inhibition. Edges represent causal relationships, where orange lines denote predicted activation and blue lines denote predicted inhibition of downstream targets. Yellow edges indicate relationships that are inconsistent with the observed expression patterns in the dataset.

Consistent with the comparison of fibrosis with control, downstream functional annotations predicted increased lipid synthesis, carbohydrate metabolism, and cellular growth-related processes, including proliferation and viability of somatic cells, indicating that metabolic reprogramming is already established at the steatosis stage. However, the upstream regulatory landscape differed with hepatic steatosis to control, with predicted activation of TP63, IL3, AHR, and XBP1, alongside inhibition of miR-16-5p and related microRNAs. These regulators are primarily associated with cellular stress responses, inflammatory signaling, and transcriptional control of protein synthesis [35], [36], [37] suggesting a broader regulatory input compared to the more lipid-centric SCAP-driven network observed in the fibrosis vs control regulator-effector networks.

At the protein level, multiple enzymes involved in synthesis and quality of carbohydrates [38], lipid biosynthesis [39], [40], and protein production [41] were interconnected, supporting coordinated upregulation of biosynthetic pathways. Notably, functions such as synthesis of protein and synthesis of carbohydrate were prominently activated in hepatic steatosis vs control, in contrast to the fibrosis vs control group where fatty acid metabolism and lipid-specific pathways were more dominant. Additionally, predicted activation of cell movement and tumor-associated processes indicates early engagement of pathways linked to cellular remodeling and adaptation.

Comparatively, while both networks characteristic to hepatic steatosis and fibrosis share activation of lipid metabolic processes, the latter network appears more specialized toward fatty acid metabolism and lipogenesis under the control of SCAP/SREBP-related mechanisms [42]. In contrast, the hepatic steatosis-control network reflects a broader, upstream-driven response integrating inflammatory cues (IL3), environmental sensing (AHR), endoplasmic reticulum stress (XBP1), and microRNA-mediated regulation [35], [43], [44]. This suggests that hepatic steatosis represents an earlier, more heterogeneous metabolic and stress-adaptive state that precedes the more focused and transcriptionally coordinated lipid metabolic reprogramming observed in fibrosis.

Overall, these findings support a model in which hepatic steatosis initiates widespread metabolic and regulatory alterations, including activation of protein and carbohydrate biosynthesis, which subsequently converge into a more defined lipid-centric metabolic phenotype during progression to fibrosis.

Canonical pathway analysis (Figure S7) further supported the regulator-effector network findings, demonstrating consistent enrichment of detoxification and metabolic pathways in both hepatic steatosis and fibrosis compared with control samples, with differences primarily in activation magnitude and pathway emphasis. Phase I (functionalization) and Phase II (conjugation) xenobiotic metabolism pathways were among the most strongly activated in both conditions, indicating enhanced hepatic detoxification and metabolic adaptation. This is in agreement with the predicted activation of stress- and environment-responsive regulators, such as AHR, observed in the network characteristic to hepatic steatosis (Figure 5A), supporting a role for xenobiotic and oxidative stress signaling in disease progression.

Consistent with the lipid metabolic reprogramming observed in the regulator-effector networks (Figure 5A, 6B), pathways related to peroxisomal function, including peroxisomal protein import and lipid metabolism, were significantly enriched, reflecting increased fatty acid handling and β-oxidation demands [45]. Similarly, enrichment of bile acid and bile salt metabolism further supports alterations in lipid and cholesterol homeostasis [46] across both conditions. The activation of retinoic acid signaling aligns with the upstream regulatory patterns identified in Figure 5A, suggesting involvement of nuclear receptor-mediated transcriptional programs linking metabolic adaptation with inflammatory and differentiation processes.

In addition, enrichment of neutrophil degranulation corroborates the inflammatory component identified in the network analysis, indicating engagement of innate immune responses [47]. Notably, fibrosis exhibited generally higher activation scores across several pathways, supporting a more pronounced and functionally focused metabolic and immune response.

Overall, these pathway-level alterations reinforce the network-based observations, indicating that hepatic steatosis is characterized by early activation of stress-responsive, detoxification, and biosynthetic pathways under a broad upstream regulatory landscape, whereas fibrosis reflects a more pronounced and transcriptionally coordinated metabolic reprogramming, particularly centered on lipid metabolism.

A direct comparison between fibrosis and hepatic steatosis using IPA canonical pathway analysis was not feasible due to the absence of robust pathway predictions. Several pathways lacked sufficient information for activity state inference, while others did not meet inclusion criteria, as fewer than four analysis-ready molecules were mapped to the respective pathways. This limitation is likely attributable to the relatively small number of differentially abundant proteins identified in this comparison (13 downregulated and 11 upregulated).

The present findings are broadly consistent with emerging proteomic evidence indicating that hepatic steatosis is characterized by widespread metabolic and regulatory reprogramming, rather than a purely lipid-centric phenotype [48], [49]. Several studies have demonstrated early activation of central carbon metabolism, protein synthesis, and endoplasmic reticulum stress pathways, including XBP1-mediated unfolded protein response signaling, supporting the notion of a stress-adaptive and biosynthetically active state in steatosis.

In agreement with this, inflammatory and cytokine-associated regulatory inputs [50], as well as microRNA-mediated derepression of proliferative pathways [51], have been reported in early disease stages, reinforcing the concept of a heterogeneous upstream regulatory landscape.

In contrast, progression to fibrosis has been associated with a more focused metabolic phenotype dominated by lipid metabolism and SCAP/SREBP-driven lipogenic signaling, suggesting a transition from a broadly adaptive to a more specialized metabolic program [52].

However, some studies report earlier activation of lipogenic pathways, indicating that lipid metabolism may already be engaged during steatosis, albeit at lower magnitude [53]. Additionally, the role of regulators such as AHR appears to be context-dependent, reflecting both adaptive and pathogenic functions [54].

Collectively, these findings support a model in which hepatic steatosis represents an early, metabolically flexible state integrating stress, inflammatory, and biosynthetic responses, which progressively converges toward a lipid-centered metabolic network during fibrosis.

## 4. Conclusion

Formaldehyde-fixed paraffin-embedded tissues represent one of the most valuable resources for clinical research. However, their application in proteomics remains limited by the lack of standardized and robust sample preparation protocols. In this study, we compared four extraction and digestion workflows for FFPE tissues, all employing xylene-free deparaffinization, with respect to proteome depth and data quality.

Our results demonstrate that a chaotropic-, reductant-, and surfactant-free in-solution digestion approach (App_2) provides high peptide and protein yield with reliable tryptic cleavage efficiency. In addition, we developed a modified version of this workflow (App_2M), which further improves peptide coverage while maintaining comparable protein depth. Notably, this protocol does not require specialized equipment, making it well suited for implementation in standard clinical laboratory settings.

To evaluate its practical applicability, the optimized workflow was applied to low-input FFPE needle biopsy samples, enabling the detection of distinct proteomic signatures across a small cohort comprising hepatic steatosis and liver fibrosis to control groups.

Although missingness remains an inherent limitation of low-input FFPE proteomics, stringent filtering criteria and exclusion of low-quality contaminated samples improved data robustness and enabled identification of biologically meaningful proteomic trends. Taken together, our findings highlight a simple, xylene-free, and MS-compatible workflow that supports efficient proteomic analysis of FFPE tissues. This approach represents a promising strategy for routine research applications and may, with further validation, be adapted for future clinical and diagnostic use.

## 5. Data and tool availability

Metadata regarding the experiments are presented in .tsv format as metadata/sdrf. The Mass Spectrometry proteomics data generated in this study have been deposited to the ProteomeXchange Consortium via the MassIVE repository under accession number PXD078226 (MassIVE accession: MSV000101787).

## 6. Supplementary Materials

- Figure S1: A global comparison of App 1-4; Figure S2: Overlap comparison of App_2 and App_3; Figure S3: Ratio of posttranslational modified peptides; Figure S4: Proportion of proteins identified by unique peptides; Figure S5: Pie chart showing intensity distribution of protein groups (PGs); Figure S6: Heatmap and principal component analysis (PCA) of study samples; Figure S7: Heatmap of the top 10 canonical pathways derived from differentially abundant proteins (DAPs) in hepatic steatosis and fibrosis (PDF)
- Table S1: DIA-NN 2.2 precursor ion generation, output, and algorithm parameters; Table S4: Liver disease case study stripped peptide sequences; Table S5: Liver disease case study DEF group-level outputs; Table S6: Differentially abundant proteins using a limma model (XLSX)
- Table S2: App_2 and App_3 DEF group-level outputs (XLSX)
- Table S3: App_2 and App_2M DEF group-level outputs (XLSX)
- Table S7: Regulator-effector network molecules and relationships (XLSX)

## 7. Author Information

## 8. Author Contributions

Conceptualization, G.M.M. and É.C.; methodology, G.M.M., L.T.; software, G.M.M.; formal analysis, G.M.M., L.B., G.M., G.K., and Z.S.; investigation, G.M.M., L.B. and G.M.; resources, É.C., L.B. and G.M.; data curation, G.M.M., G.K., and Z.S.; writing—original draft preparation, G.M.M., G.M., and G.K.; writing—review and editing, É.C., Z.S., L.T. and G.M.; visualization, G.M.M.; supervision, É.C.; project administration, É.C.; funding acquisition, É.C., Z.S., and L.T. All authors have read and agreed to the published version of the manuscript.

## 10. Institutional Review Board Statement

The study was conducted in accordance with the Declaration of Helsinki, and approved by the Ethics Committee of the University of Debrecen (IV/8465-3/2021/EKU).

## 11. Acknowledgments

The authors sincerely thank Dr. Christine Carapito and Dr. Jeewan Babu Rijal for hosting the research visit and for their valuable scientific discussions, technical support, and access to laboratory infrastructure. We also thank the help of Dr. Gergő Kalló for the critical review of the manuscript.

## 12. Abbreviations

LC-MS/MS: Liquid chromatography-tandem mass spectrometry
FFPE: Formaldehyde-fixed paraffin-embedded
LCM: Laser capture microdissection
MMD: Manual microdissection
AFA: Adaptive focused acoustics
PHAD: Projected hot air deparaffinization
SDS: Sodium dodecyl sulfate
SDC: Sodium deoxycholate
App: Approach
HIAR: Heat-induced antigen retrieval
DTT: Dithiothreitol
TFA: Trifluoroacetic acid
ACN: Acetonitrile
TEAB: Triethylammonium bicarbonate
ABC: Ammonium bicarbonate
FA: Formic acid
NSI: Nanoelectrospray ion source
iRT: Indexed retention time
PCA: Principal component analysis
FDR: False discovery rate
DAPs: Differentially abundant proteins
IPA: Ingenuity pathway analysis.

